# Low and high frequency signatures of impaired consciousness in temporal lobe seizures

**DOI:** 10.1101/2025.07.01.662627

**Authors:** Taruna Yadav, Bogdan Patedakis Litvinov, George W. Culler, Avisha Kumar, Aya Khalaf, Yinchen Song, Benjamin H. Brinkmann, Boney Joseph, Zan Ahmad, Nisali Gunawardane, Violeta Contreras Ramirez, Imran H Quraishi, START Clinical Trial Research Consortium, Barbara C. Jobst, Gregory Worrell, Hal Blumenfeld

## Abstract

Impaired consciousness is a debilitating and unpredictable outcome of mesial temporal lobe seizures whose mechanisms to date remain unclear. Moreover, questions about the relationship between impaired consciousness and lateralization, hemispheric spread and electrophysiological characteristics of seizures are yet to be answered. To address these gaps, we conducted in-depth investigation of behavioral and intracranial EEG data from 186 mesial temporal lobe seizures of 51 patients with intractable mesial temporal lobe epilepsy. We found that bilateral mesial temporal spread of seizures is not a necessary condition for impaired consciousness, although seizures with bilateral mesial temporal involvement were significantly more likely to have impaired consciousness than unilateral seizures. Contrary to prior belief, we found no relationship between the onset side (left vs. right temporal lobe or language dominant vs. non-dominant lobe) of seizures and the probability of impaired consciousness. Lastly, we established that widespread increases in slow-wave activity (delta band) in extratemporal cortical areas, as well as increases in fast activity (beta band) in the temporal lobes were both robust markers of seizures with impaired consciousness and could predict ictal impairment with up to 86% accuracy. Our findings shed new light on networks that underlie impaired consciousness in temporal lobe epilepsy and may help guide deep brain stimulation of such systems (e.g. via thalamic nuclei) as a potential intervention to improve consciousness during seizures.

## Introduction

Consciousness is an umbrella term with different definitions across fields such as philosophy, psychology and, medical science. In clinical research, consciousness is understood as a continuum with varying levels of arousal or alertness and awareness extending from normal wakefulness to deep sleep to coma or vegetative state (Gosseries et al., 2011). Disorders of consciousness could be observed following an injury (e.g. traumatic brain injury) or due to neurological disorders or diseases (e.g. stroke and epilepsy) (Gosseries et al., 2011). Temporal lobe epilepsy (TLE) is the most common type of focal epilepsy originating in the limbic structures such as hippocampus and amygdala. Impaired consciousness is often a serious manifestation of TLE seizures negatively affecting quality of life and safety of patients. (Téllez-Zenteno & Hernández-Ronquillo, 2012). Studies of the mechanisms underlying impaired consciousness during TLE seizures suggest the involvement of cortical and sub-cortical structures. Intracranial EEG recordings reveal correlation between increased synchrony between the temporal lobe and thalamus during the early and late seizure periods of mesial temporal lobe seizures correlates and early loss of consciousness (Guye et al., 2006). Moreover, excessive and prolonged long distance cortico-cortical and thalamo-cortical synchronization involving extra-temporal structures such as parietal cortices is strongly linked to impaired consciousness (Arthuis et al., 2009). An alternate hypothesis (network-inhibition hypothesis) of underlying mechanism emphasizes that propagation of ictal discharges beyond the temporal lobe may not be necessary and simultaneous irregular neocortical slowing may be responsible for impairment (Blumenfeld, Rivera, et al., 2004). This hypothesis suggest that abnormal activation of upper brain stem and mesial diencephalon and wide-spread abnormal inhibition of frontal and parietal cortices could be responsible for impaired consciousness (Blumenfeld, McNally, et al., 2004; Englot et al., 2010). Evidence from an awake mouse model of focal limbic seizures shows that impaired responsiveness to auditory stimuli is related to increased slow wave activity and altered acetylcholine neurotransmission in the orbital frontal cortex (Sieu et al., 2024) and decreased multi-unit activity in locus coeruleus (Valcarce-Aspegren et al., 2025). While these studies shed light on possible mechanisms underlying impaired consciousness, they are limited by the sample size, number of electrodes and hemispheric coverage. Moreover, whether the observations about csignificant cortical slowing and limbic fast activity have utility in predicting impairment at single seizure level hasn’t been tested before.

With this study, we tackle the limitations of the previous studies and address three long standing questions about the relationship between clinically relevant seizure characteristics through comprehensive analysis of large multi-center intracranial EEG datasets. We asked, i) are bilateral seizures more likely to cause impairment than unilateral seizures, ii) are left sided seizures more probable to cause impairment than right sided seizures, and iii) is extra-temporal slow wave activity predictive of impairment during seizures? We additionally report distinct frequency band specific changes in nine brain regions of interest involving mesial temporal, lateral temporal and seven other extra-temporal regions. As per the latest ILAE seizure classification, we labeled seizures as focal impaired consciousness (FIC) or focal preserved consciousness (FPC) seizures based on evaluation of ictal responsiveness and postictal recall behavior (Beniczky et al., 2025) .

## Methods

### Patients

All procedures were in accordance with the institutional review boards for human studies at the four data collection sites (Yale University School of Medicine, Mayo Clinic, Dartmouth-Hitchcock medical center and New York University School of Medicine), as well as National Institutes of Health guidelines. Informed consent was obtained from all subjects.

Inclusion and exclusion criteria were chosen to identify a homogenous group of patients with confirmed mesial temporal lobe epilepsy who had undergone intracranial EEG and video monitoring. Patients with intracranial recordings between 1995-2008 and 2016-2019 at Yale University, between 2003-2004 at NYU, between 2013-2021 at Mayo Clinic and between 2015-2020 at Dartmouth medical center were included in the analysis. Patients with mesial temporal onset based on intracranial EEG were included in the analysis. The following seizures were excluded from analysis: focal to bilateral tonic-clonic seizures and seizures in which intact or impaired behavioral responsiveness could not be determined (e.g. no one interacted with the patient during the seizure or video not available). A total of 186 seizures from 51 patients (23 female, mean age 38 yrs (range 17-69 years)) were used for the analysis (see Figure S 1 for consort diagram summarizing seizure exclusion criteria). Forty-one patients were right-handed, eight were left-handed and two were ambidextrous. Forty-two patients underwent Wada testing via the intracarotid sodium amobarbital procedure as part of a preoperative neuropsychological testing battery. Thirty-eight patients were found to have left hemispheric dominance for language production while four had right dominance.

### Anatomic localization of electrode positions

Surgical implants included subdural strip, grid and depth electrode contacts (AdTech Medical Instruments Corp., Racine, WI, USA for Yale-Englot and Dartmouth, DIXI Microdeep electrodes, DIXI medical, Chaudefontaine, France and sEEG Depthalon, PMT Corporation, Chanhassen, MN for Mayo).

Preoperative planning of intracranial electrode locations was decided in each case based on clinical grounds; therefore, electrode positions were not standardized. However, electrode coverage in all cases included at least one temporal lobe and several frontal or parietal neocortical contacts ipsilateral and/or contralateral to the side of seizure onset. Bilateral electrodes were implanted in 25 of the 51 patients. Depth electrodes, together with strip electrodes, were used to study mesial and lateral temporal structures, while grids and strips were most used for other regions such as lateral fronto-parietal-temporal cortex.

High-resolution MRI scans were performed on all patients after intracranial electrode implantation using 3D volume inversion recovery prepped fast spoiled gradient recalled echo (IR-FSPGR) imaging on a 1.5 T and 3T systems at Yale and at 1.5 T at NYU, 3D T1-weigted gradient-echo sequence (magnetization-prepared rapid gradient echo, MPRAGE) imaging on a 3-T MR scanner (Philips and Siemens) at Dartmouth and using variable intensity scanners (7T Siemens Terra, 3T Siemens Verio, 3T GE PET/MR Signa or 1.5T GE)) at Mayo. High-resolution CT images were obtained after the intracranial electrode implantation and co-registered with prior T1 images to localize the electrode positions in each patient.

The brain surface was segmented into the following nine anatomical regions for both ipsilateral and contralateral hemispheres: mesial temporal, lateral temporal, orbital frontal, lateral frontal (includes pre-central gyri), medial frontal, lateral parietal (includes post-central gyri), medial parietal, insula and occipital. Boundaries used for these anatomic regions have been shown and described in detail previously (Blumenfeld, Rivera, et al., 2004). Electrode contacts identified on the MRI scans were assigned to these regions for EEG analysis, as previously described (Blumenfeld, Rivera, et al., 2004). Contacts lying intermediate between two anatomic regions or in the subcortical white matter were excluded from analysis. A total of 4586 grey matter electrodes (3427 ipsilateral and 1159 contralateral to seizure onset side) were available for EEG analysis from 51 patients.

Electrode locations in MNI space (Colin 22 brain template) were identified based on pre-op MRI and post-op CT using BioImage Suite (http://bioimagesuite.yale.edu/) for patients whose MRI and CT scans were available. A triangular mesh representing the standard MNI cortical surface was created in BioImage Suite and electrodes were assigned to the nearest vertex. For visualization, we used the MNI2FS MATLAB toolbox to render MNI coordinates on a canonical FreeSurfer inflated brain mesh (Price, 2017) at level 5 inflation.

### Intracranial electroencephalography recordings

For patients at Yale (1998-2008), intracranial EEG (icEEG) signals were recorded continuously using Telefactor Beehive systems (Grass Telefactor, Astro-Med, Inc., West Warwick, RI, USA) or Bio-Logic Systems 128-channel clinical EEG and video monitoring equipment (Bio-Logic Systems Corp., Mundelein, IL, USA), sampled at 200 Hz. For patients at Yale (2016-2019), signals were recorded at 1024 Hz sampling rate with Natus Neurolink 256-channel amplifier or between 512-4096 Hz sampling rate for Natus Quantum 256-channel amplifier (Natus, Middleton, WI). For patients at New York University, 128 icEEG channels were acquired using the BMSI 5000/6000 EEG system (Nicolet Biomedical, Inc., Madison, WI, USA) at 400 Hz sampling rate. For patients at Dartmouth medical center, icEEG signals were recorded using GRASS TWin EEG system (Grass Technologies Corp.) or Quantum 256-channel amplifier at 400 Hz sampling rate for GRASS system or 2048 Hz for Natus system. For patients at Mayo clinic, icEEG signals were obtained using Quantum 256-channel amplifier at 500 Hz sampling rate.

### Electroencephalography analysis

Prior to analysis, icEEG recordings were manually reviewed to identify seizure onset and offset. Seizure onset and offset were defined based on the onset or termination of poly-spike activity or low-voltage fast activity in the frequency range 10-45 Hz. Seizure onset was specifically identified from mesial temporal contacts of icEEG, but seizure offset could be defined based on the contacts outside of the mesial temporal lobe, depending on the spread of seizure.

EEG records were exported from 60 sec before seizure onset and 60 sec after the seizure offset. Artefacts were delineated on the record based on visual inspection. For further analysis, EEGs were aligned to seizure onset and seizure offset to extract a 30-sec baseline, up to 300 sec ictal and 70 sec postictal periods. Channels with artefacts throughout the EEG data were rejected.

Remaining channels were inspected for artifacts in 1 sec segments and those with artefacts were removed from analysis. Next, a band stop filter (2^nd^ order Butterworth 59-61 Hz) was applied to remove electrical noise from EEG signals.

For each electrode, power spectral density (PSD) estimate was computed by applying fast Fourier transform (FFT) with hamming window on 1-sec non-overlapping EEG segments at a frequency resolution of 1 Hz (FFT size = EEG segment size). Next, band power was calculated from PSD by averaging the power corresponding to each of the following frequency bands: delta (1 to 4 Hz), theta (5 to 8 Hz), alpha (9–13 Hz), beta (14 to 25 Hz) and gamma (26–50 Hz). EEG power was quantified as percent change from mean baseline [100 x (EEG signal power – mean baseline power)/ mean baseline power]. Next, computed EEG power from each seizure were further averaged into 10-sec non-overlapping bins. Each seizure’s band EEG power time-courses were checked for outliers. Ictal and postictal bins with power value outside 4 SD from mean ictal and postictal powers respectively were considered outliers and removed from analysis.

Statistical group analyses were conducted using n-way ANOVA model with interaction effects grouped by four variables namely, seizure type (FIC or FPC), brain region (nine anatomical regions of interest defined previously), frequency band (Delta, Theta, Alpha, Beta or Gamma) and seizure period of interest (ictal or postictal). For variables with significant coefficients in ANOVA, post-hoc pairwise tests were conducted using non-parametric Wilcoxon Ranksum test and resulting p-values were corrected for multiple comparisons using Benjamini-Hochberg False Discovery Rate method. ANOVA models were run separately for ipsilateral and contralateral electrodes. All statistical tests were performed using MATLAB 2020a (MathWorks, Natick, MA, USA) with significance assessed at p <= 0.05. Due to a lack of alternative non-parametric test and known robustness against deviations from normal distribution of data, we think the use of ANOVA model was appropriate for initial group level statistical testing.

### Brain map generation

Brain maps were generated using BrainNet viewer, a MATLAB based brain visualization toolbox (Xia, Wang, & He, 2013). Standard ICBM152 brain surface in MNI space was used. WFU (Wake Forest University School of Medicine) pick atlas tool (Maldjian, Laurienti, & Burdette, 2004; Maldjian, Laurienti, Kraft, & Burdette, 2003; Tzourio-Mazoyer et al., 2002) was next used to load an atlas (AAL MNI v4) and map the regions of interest to the brain surface.

Boundaries for the nine regions of interest in this work were identified based on the atlas and manually traced for later editing. Next, mean percent change in ictal and postictal powers of each brain region were natural log transformed, assigned a color code and finally edited in Adobe Illustrator.

### Behavioral analysis

Videos of all seizures with onset in mesial temporal lobe were viewed and rated on a scale of 1-4 by a reviewer who was provided with seizure onset and offset times based on intracranial EEG. All patients included in the study were presented with questions or commands during seizures by either medical staff or family members. Stimuli such loud sounds or conversations not involving the patient that elicit orienting behaviors were not considered towards assessment. Focal to bilateral tonic-clonic seizures, and seizures with no video available for behavioral assessment were excluded from the analysis. Seizures in which staff or family members did not interact with the patient at any time during the seizure were rated as 1 and excluded from further analysis.

Seizures where interaction happened but the patient’s response was unclear were rated with score of 2. Seizures in which patients had any impaired responsiveness to verbal questions, failed to correctly follow simple commands at any point during the seizure or exhibited amnesia were classified as focal impaired consciousness (FIC) seizures and rated with score 3. Seizures where patients remained fully alert and interacted appropriately throughout the event were considered focal preserved consciousness (FPC) seizures and rated with score 4. Only seizures with ratings 3 and 4 were included in electrophysiological analysis.

## Results

We studied intracranial EEG recordings during 96 FPC (or spared) and 90 FIC (or impaired) seizures in 51 patients, with behavioral seizure type determined by responsiveness ratings observed on video-EEG. FIC seizures were significantly longer than FPC seizures mean (±SEM) duration, 131.1 (±13.8) s and 94.8 (±9.7) s respectively (p = 3.19 x 10^-9^, Wilcoxon Ranksum test).

### Greater extratemporal slow and temporal lobe fast activity in FIC seizures

To examine the differences in electrographic characteristics of seizures with or without impaired consciousness, we analyzed icEEG from ipsilateral and contralateral contacts during up to 370 s ictal and 60 s postictal period. Typical progression of a FPC and FIC seizure is shown in Figure 1. Sample icEEG recordings show seizure activity during onset, early and late ictal and postictal periods. Both FPC and FIC seizures had similar onset pattern consisting of low-voltage fast activity in the mesial temporal lobe followed by poly-spike and sharp wave patterns in the mesial and lateral temporal lobe during the early and late ictal periods. Moreover, during the early period, FIC seizures displayed an asynchronous and irregular slow wave activity in the lateral temporal and frontoparietal regions which continued in the later ictal and postictal periods and spread to various extra-temporal regions (Figure 1B), which was not observed in FPC seizures. Figure 2 shows band power time courses and bar plots for FPC and FIC seizures during the ictal (first 150 s) and postictal periods (60 s). Data present percentage change in icEEG power relative to baseline (30 s period before seizure onset) in 10 second bins from 180 seizures of 49 patients. For each seizure, icEEG power was aggregated from the mesial and lateral temporal contacts to compute mean time courses for the temporal region to generate ipsilateral Temporal (Figure 2 A, E) and contralateral Temporal (Figure 2 C, G) plots. Similarly, electrode contacts from all other brain regions were combined across seizures to generate ipsilateral Extra-Temporal (Figure 2 B, F) and contralateral Extra-Temporal plots (Figure 2 D, H). Bar plots (Figure 2 I though L) show the mean percent change in ictal and postictal power. In the ipsilateral temporal region, icEEG power of FIC seizures was significantly greater than that of FPC seizures in the ictal period in all the frequency bands with largest increase in the beta band. For FPC seizures, broadband increase in ictal power was observed on the ipsilateral temporal regions but such changes were minimally observed on the contralateral temporal regions. FIC seizures on the other hand had broadband increase in both ipsilateral and contralateral temporal regions but the changes were much larger on ipsilateral side. Changes in icEEG power during the postictal period in the temporal lobes were relatively small in magnitude for both FPC and FIC seizures. Ictal icEEG power increases in extra-temporal regions were dominated by delta band both on ipsilateral and contralateral sides in addition to changes in other frequency bands. FIC seizures had significantly larger bilateral slow wave activity (delta band power) than FPC seizures both during the ictal and postictal periods. Additional time course and bar plots for each of nine brain regions are presented in the supplementary section (Figure S 3, Figure S 4, Figure S 5).

**Figure 1.**
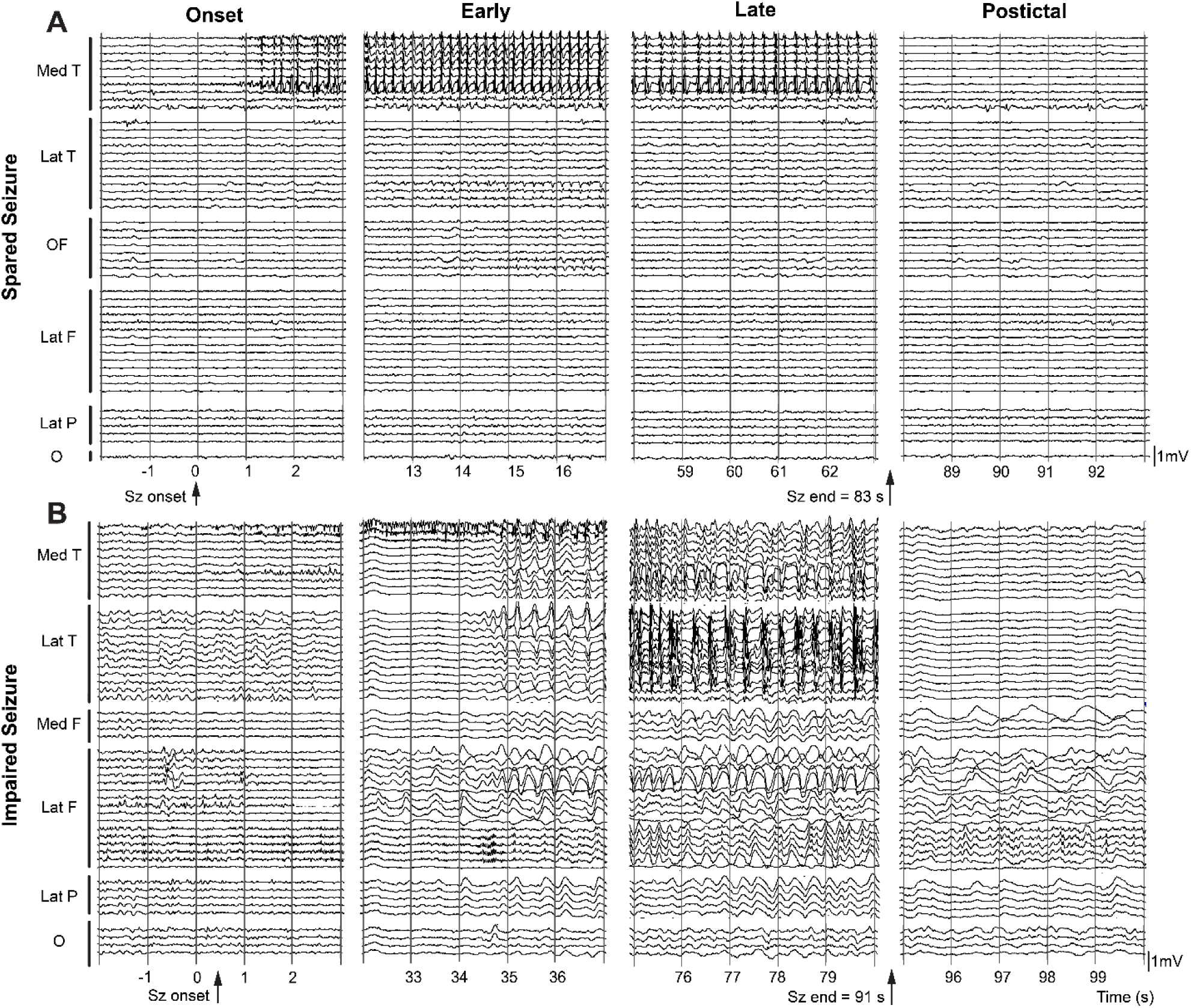
Example intracranial EEG recordings during a sample focal preserved consciousness (FPC, spared) and focal impaired consciousness (FIC, impaired) temporal lobe seizure. EEG is shown for ipsilateral contacts for four seizure periods (onset, early, late and postictal) from different brain regions. (A) Spared seizure onset is seen as low-voltage fast activity on mesial temporal contacts which evolves to rhythmic poly-spike and sharp wave activity (early and late periods) and largely remains in the mesial temporal lobe. Postictal suppression is seen in the mesial temporal lobe with no large-amplitude neocortical slow activity. (B) Impaired seizure onset has similar low-voltage fast pattern followed by rhythmic poly-spike and sharp wave activity in the mesial and lateral temporal contacts during the early and late periods and is suppressed during the postictal period. Frontal and parietal contacts show irregular slow wave activity asynchronous to fast activity in mesial temporal lobe. Large amplitude neocortical slowing continues throughout the early and later periods and persists in the postictal period. Mes T = mesial temporal; Lat T = lateral temporal; OF= orbital frontal; Lat F = lateral frontal; Lat P = lateral parietal; O= occipital.

**Figure 2.**
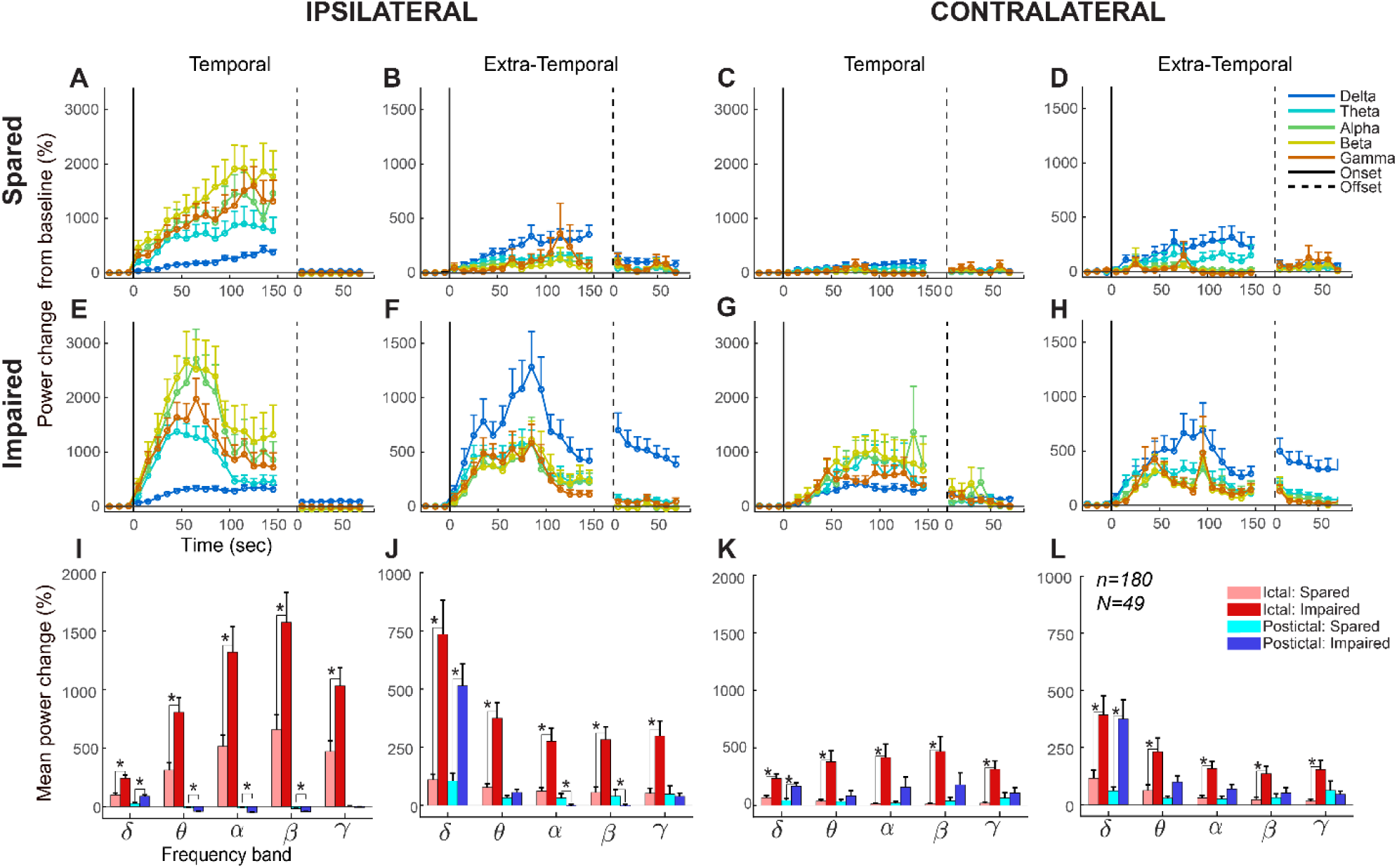
Time course plots of intracranial EEG power changes during spared and impaired seizures (A) & (B) Ipsilateral to onset of spared seizures for temporal and extra-temporal electrodes respectively. (C) & (D) Contralateral to onset of spared seizures for temporal and extra-temporal brain regions respectively. (E), (F), (G) and (H) show similar ipsilateral and contralateral temporal and extra-temporal electrodes for impaired seizures. Data are mean percent change (+SEM) in EEG power from 30 s pre-seizure baseline to 150 s ictal and 60 s postictal period binned every 10 s. Vertical solid lines indicate seizure onset and dotted lines indicate seizure offset. (I), (J), (K) and (L) show mean (+ SEM) percent change in power during ictal (pink vs. red) and postictal (cyan vs. blue) periods for spared and impaired seizures respectively in the different frequency bands (Delta, Theta, Alpha, Beta and Gamma). Note: different scale between temporal and extra-temporal data plots. Impaired seizures show significantly (p<0.05, after correction for multiple comparisons) larger broadband increases compared to spared seizures in the ipsilateral temporal and extra-temporal lobes with highest increases in the beta and delta bands respectively. Postictally, temporal lobe shows broadband suppression while extra-temporal lobe shows sustained slow-wave activity (delta band power) for impaired seizures. Similar patterns were observed for contralateral temporal and extra-temporal lobes but with comparatively lower magnitudes.

For visual comparison of icEEG power between FPC and FIC seizures, we generated brain maps summarizing power changes across nine brain regions (Figure 3). Brain maps show natural log transformed mean percentage change in power in each brain region during the ictal and postictal periods. Only brain regions with percentage change in power above a set threshold (100% for delta band and 200% for beta band) are displayed. Results are shown for beta (Figure 3 A-D) band in the ictal period and delta band in the ictal (Figure 3 E-H) and postictal (Figure 3 I-L) periods. Impaired seizures compared to spared seizures show similar beta band power in the ipsilateral mesial temporal lobe but much higher power in the ipsilateral lateral temporal, mesial frontal, orbitofrontal and lateral parietal lobes, and contralateral mesial temporal lobe ((A) and (C)). Beta increase in impaired seizures is restricted to contralateral mesial temporal lobe with no contralateral activity for spared seizures ((B) and (D)). In the ictal period, impaired seizures show bilateral widespread cortical slowing (delta band power increases) while delta band increases are only observed in the ipsilateral mesial and contralateral lateral frontal regions for spared seizures (E) and (F)). Bilateral neocortical slow wave activity sustained postictally in the FIC seizures but not for FPC seizures.

**Figure 3.**
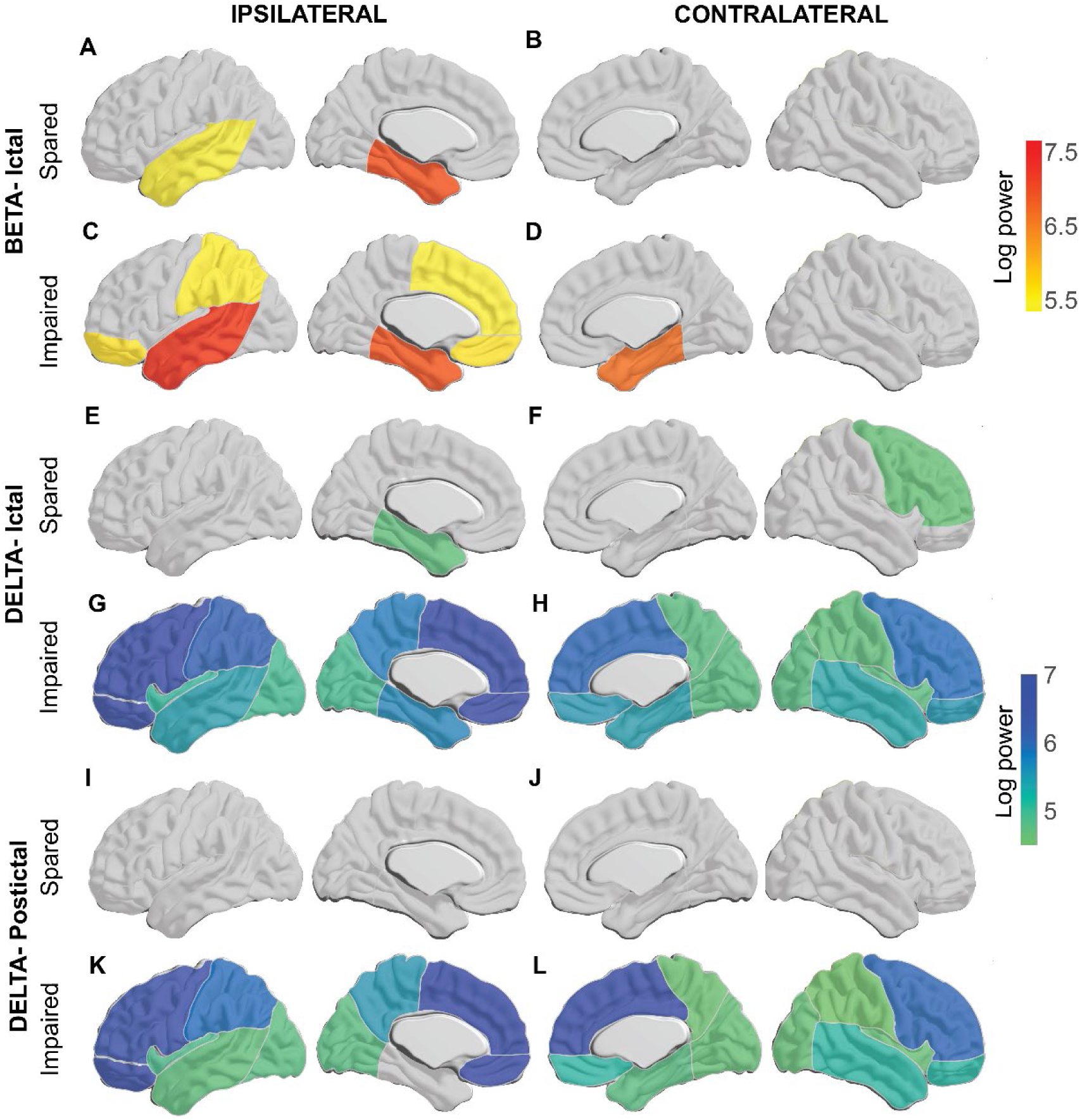
Brain maps show mean percentage change in power from baseline in the beta band (warm colors) during ictal period (A through D) and delta band (cool colors) during the ictal (E though H) and postictal periods (I through L) for spared and impaired seizures. Mean power from nine brain regions (mesial temporal, lateral temporal, mesial frontal, lateral frontal, orbitofrontal, mesial parietal, lateral parietal, insula, and occipital) were converted to natural log scale for visualization. A threshold of 200% and 100% change in mean power was applied for beta and delta bands respectively and only regions with change>= threshold are shown. Impaired seizures compared to spared seizures show similar beta band power in the ipsilateral mesial temporal lobe but much higher power in the lateral temporal, mesial frontal, orbitofrontal and lateral parietal lobes ((A) and (C)). Beta increase in impaired seizures is restricted to contralateral mesial temporal lobe with no contralateral activity for spared seizures ((B) and (D)). In the ictal period, impaired seizures show bilateral widespread cortical slowing (delta band power increases) while delta band increases are only observed in the ipsilateral mesial and contralateral lateral frontal regions for spared seizures ((E) and (F)). Postictally, bilateral neocortical slow wave activity sustains in the impaired seizures but no delta band increase are sustained for spared seizures.

### Effect of seizure lateralization and onset side on seizure severity

To answer the long-standing questions about the relationship between seizure spread and onset side with probability of impaired consciousness during seizures, we performed the following analysis. Seizures with bilateral implants in the mesial temporal lobe (MTL) were chosen and segregated based on seizure spread. icEEG beta power from ipsilateral and contralateral mesial temporal contacts were visually reviewed to determine if a seizure was unilateral (seizure started on the ipsilateral MTL remained on that side throughout the seizure duration) or bilateral (seizure started simultaneously on both MTL or seizure started on ipsilateral MTL and spread to the contralateral MTL at any point during the ictal period). Of 29 bilateral seizures, 24 seizures had impaired consciousness while of 57 unilateral seizures, 21 seizures had impaired consciousness (Figure 4 A). These results show that bilateral seizures are significantly more likely to cause impairment than unilateral seizures (χ^2^= 16.2, p<0.001). It must be noted though that spread of seizure activity to contralateral MTL is not necessary for impairment. Unilateral seizures, despite their lower likelihood compared to bilateral seizures can cause impaired consciousness as observed in 37% of unilateral seizures here.

**Figure 4.**
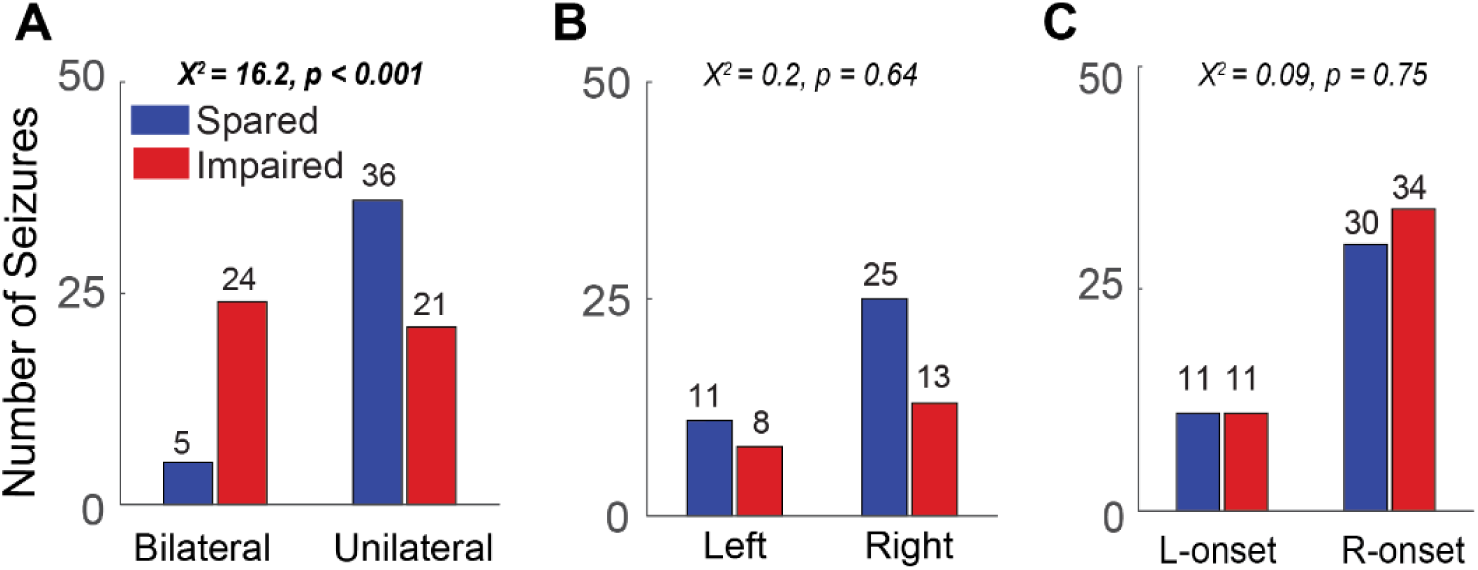
Histograms showing seizure count in different categories. Note: Seizures with bilateral involvement are called bilateral seizures henceforth, seizures that remain on one side are called unilateral seizures. (A) Unilateral vs. bilateral seizures verified by bilateral recordings, (B) verified unilateral seizures grouped by type (left-only vs. right-only), (C) verified unilateral and bilateral seizures grouped by onset side (left vs. right). Chi-square statistic with p-value in each plot shows if the number of spared and impaired seizures varied significantly with the grouping variable (x-axis).

Next, for all seizures with bilateral mesial temporal implants, we lateralized the seizures based on either: 1. the temporal lobe side involved throughout the seizure, excluding any seizures with contralateral temporal lobe spread (Left vs. Right seizures; Figure 4 B); or 2. the side of temporal lobe seizure onset, even if the seizure later spread to the other side (Left-onset vs. Right-onset seizures; Figure 4 C). Of 57 unilateral seizures, 19 were left-sided and 38 were right-sided. Both left-sided and right-sided seizures could lead to impaired consciousness, with left-sided seizures not significantly more likely to cause impairment than right-sided seizures (χ^2^= 0.2, p=0.64).

Next, we considered all seizures with bilateral implants in MTL, including seizures with spread to the contralateral side. Seizures with onset on either the left or right side could lead to impaired consciousness with nearly equal probability. No side had significantly higher likelihood of causing seizures with impaired consciousness (χ^2^= 0.09, p=0.75). We also investigated the distribution of seizures for additional categories (onset on language dominant vs. non-dominant hemisphere for unilateral-only, bilateral-only and all implants) which are summarized in the supplementary section (Figure S 6).

### Predictive performance of Beta and Delta band icEEG power

Results in Figure 2 and Figure S 4 indicate that FIC seizures have largest significant differences in the mesial temporal (MT) and lateral temporal (LT) beta band power (ictal) and extra-temporal (ET) delta band power (ictal and postictal) compared to FPC seizures. To test whether these icEEG power features (MT Beta, LT Beta and ET Delta) can predict the severity of seizure, we used two approaches, i) use a machine learning model to classify a seizure into FIC or FPC and, ii) create ROC curves for each feature at different threshold power levels to determine the strength of that feature in distinguishing an FPC seizure from FIC seizure. For approach i), a support vector machine (SVM) was trained on either ipsilateral-only icEEG features or features from both ipsilateral and contralateral sides using 10-fold cross validation and repeated over 10 iterations. Classification accuracy was defined as the sum of true positives (TP) and true negatives (TN) divided by the total number of seizures. Figure 5 shows the classification accuracy when using ipsilateral-only features vs. both ipsilateral and contralateral features from the ictal (top row) and postictal periods (bottom row). For ipsilateral features, results are shown for all possible feature vectors; for bilateral features, results are only shown for top 12 feature vector among 63 possible feature vectors. Ictally, for ipsilateral features only, best accuracy was achieved for ipsilateral ET Delta (77%). For a combination of ipsilateral and contralateral features, best accuracy was achieved for ipsilateral MT beta, LT beta and ET delta together with contralateral LT beta (86%). Ictal accuracy decreased rapidly when another ipsilateral feature was added to feature vector while the accuracy varied slightly with addition of new features to the bilateral feature vector. Ictally, highest accuracy from a single predictor could be achieved with ipsilateral ET Delta (77%) or contralateral ET Delta (80%; when available). In the postictal period, classification accuracy remained consistently good with different ipsilateral features but the best performance was achieved for ipsilateral ET Delta and LT Beta (80%) and for bilateral ET Delta and LT Beta (78%; when available). Postictally, ipsilateral LT beta was the strongest single predictor of impairment with accuracy of 79% while it was ipsilateral ET Delta when bilateral features were available (75%). Overall, these results suggest that extra-temporal delta (neocortical slowing) and lateral temporal beta power are strong predictors of impairment in the ictal and postictal periods.

**Figure 5.**
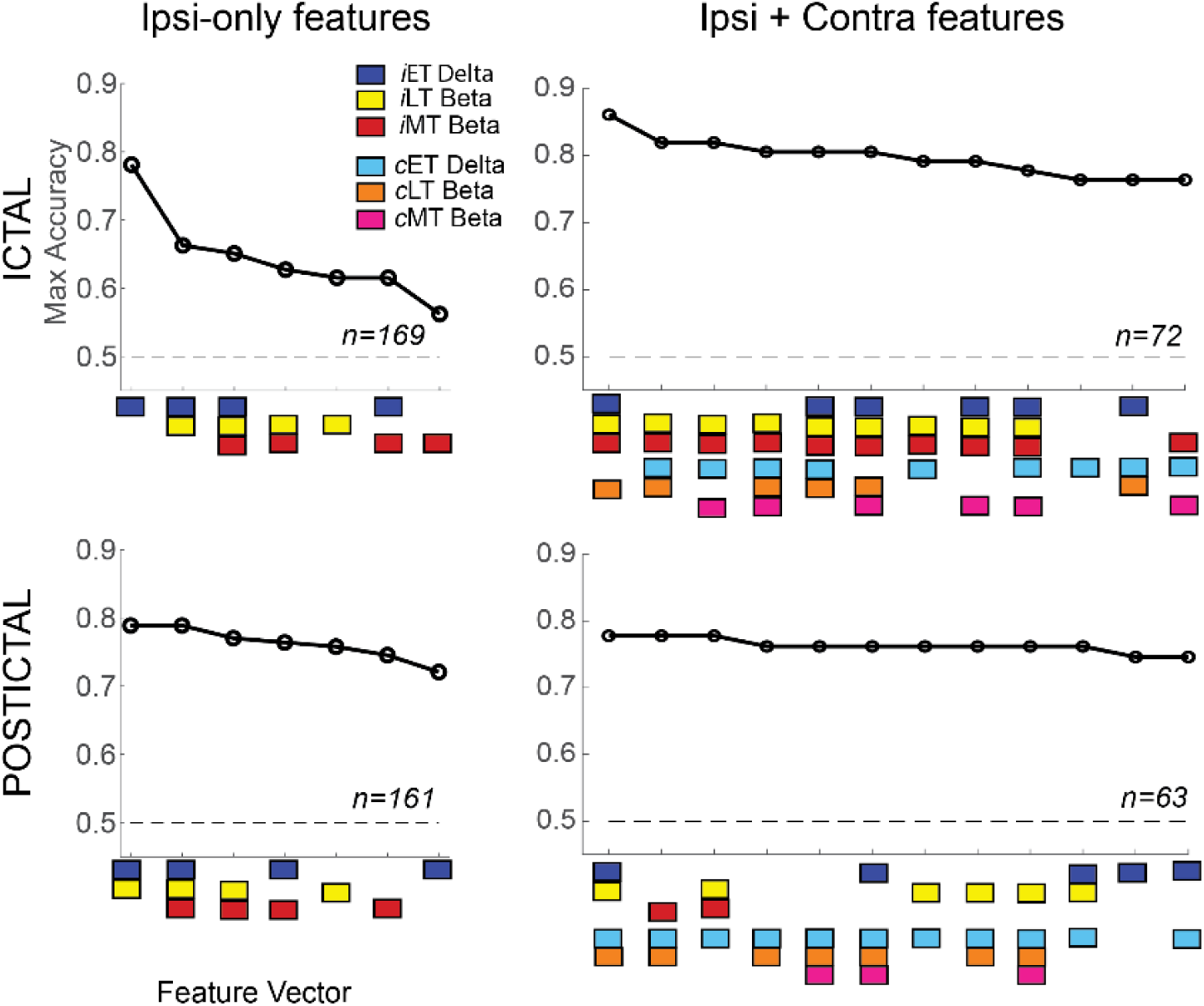
Classification accuracy (impaired vs. spared) of an SVM model trained on ipsilateral-only and ipsilateral and contralateral features for ictal and postictal periods. Features used: mean extra-temporal (ET) delta power, mean lateral temporal (LT) and mesial temporal (MT) beta power. Feature vectors (x-axis) are color-coded and rank-ordered based on maximum classification accuracy (among 10 iterations of 10-fold cross validation). Best accuracy of 77% and 86% were achieved ictally and 80% and 77% postictally using ipsilateral-only and bilateral features respectively. Ipsilateral-only ET delta and contralateral ET delta were best single predictors in the ictal period. Postictally, ipsilateral-only LT beta and ET delta achieved similar accuracy while a variety of bilateral feature combinations achieved comparable accuracies.

Next, to generate ROC curves (Figure 6), we defined sensitivity and specificity based on different chosen threshold levels for icEEG power. Threshold value represented the mean percentage change in power from baseline during the period of interest (ictal or postictal) and for chosen feature (MT beta, LT beta and ET delta). For each feature and at each threshold, we counted FIC seizures with power > threshold (considered True Positive; TP) and power <= threshold (considered False Negative; FN). Similarly, we counted FPC seizures with power> threshold (considered False Positive; FP) and power<=threshold (considered True Positive; TP).

**Figure 6.**
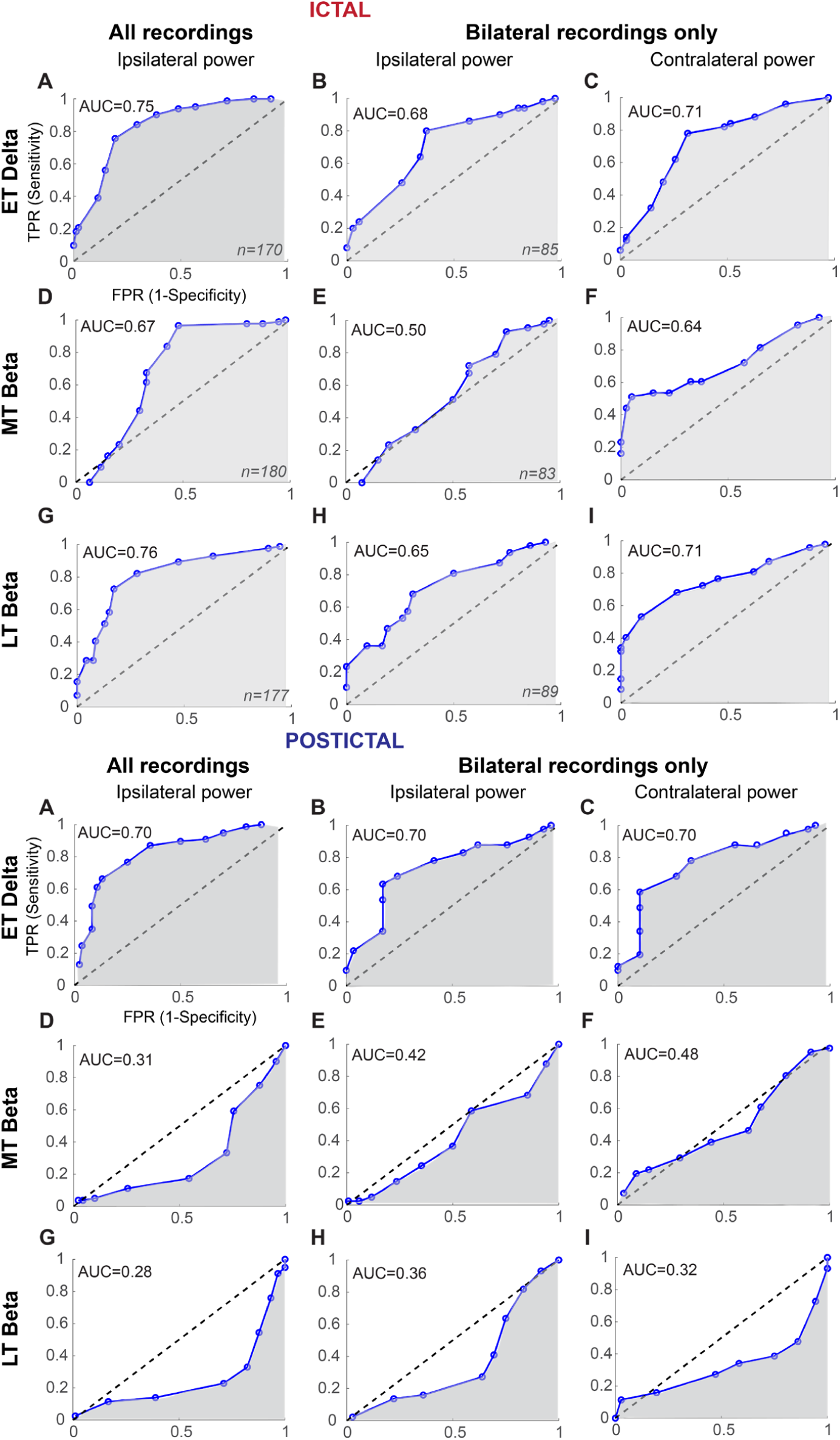
Receiver operating characteristic (ROC) and area under the curve (AUC) compare performance of different features (mean ET delta, MT beta and LT beta powers) at varying thresholds (each circle on the ROC) in the ictal (top) and postictal (bottom) plots. AUC=0.5 (dotted diagonal) represents random classification and AUC closer to 1 indicates that the feature could predict with higher probability if a seizure was impaired or spared. Ictally, both ET delta and LT beta had similar AUC for both ipsilateral and contralateral features and could be considered good predictors for impaired seizures. Postictally, both ipsilateral and contralateral ET delta achieved good AUC (0.7). True Positive Rate (TPR) = number of impaired seizures with > threshold power / number of impaired seizures, False Positive Rate (FPR)= number of spared seizures with >threshold power / number of spared seizures.

True Positive Rate (TPR, Y-axis) was defined as TP/(TP+FN) and False Positive Rate (FPR, X-axis) was defined as FP/(FP+TN). Plots (Figure 6 A, D and G) show ROC curve obtained from all seizures (i.e. seizures with ipsilateral implants) while plots (B-C, E-F, H-I) were obtained from seizures with bilateral implants in the respective brain region (mesial temporal, lateral temporal and extra-temporal). For seizures with bilateral implants, ROC curves were separately generated for ipsilateral and contralateral features. The area under the curve (AUC) was computed as ∑_𝑖_ (*TPR*(𝑖 + 1) + *TPR*(𝑖)) ∗ (*FPR*(𝑖 + 1) − *FPR*(𝑖))⁄2 where *i* represents a threshold value at which TPR and FPR are computed. In the ictal period, we observed the highest AUC for ipsilateral implants with comparable AUC for ET Delta (AUC= 0.75) and LT Beta (AUC= 0.76) i.e. these features could individually discriminate between FIC and FPC seizures with 75-76% accuracy. For seizures with bilateral implants, the best discrimination (AUC= 0.71) was achieved with contralateral ET Delta and contralateral LT Beta. AUC for MT Beta was lowest in each case (Ictal Figure 6 D, E and F) and thus was not as strong predictor of impairment in the ictal period as the other two features. AUC closer to 0.5 meant the discrimination performance was only as good as a random classifier. Overall, in the ictal period, power in the slow-wave activity in the extra-temporal region and seizure activity in the lateral temporal region (both ipsilateral and contralateral, when available) could discriminate whether a seizure would preserve or impair consciousness. In the postictal period, only ET Delta could distinguish FIC from FPC with AUC= 0.70. Both MT beta and LT beta had poor discriminative performance with (AUC <=0.5). This indicates after the seizure has ended, activity in the mesial or lateral temporal regions (either ipsilateral or contralateral) couldn’t discriminate if the seizure preserved or impaired consciousness. Only neocortical slowing in the extra-temporal regions (similar performance with ipsilateral or contralateral feature) could discriminate between FIC and FPC seizures with 70% accuracy.

## Discussion

In this work, we employed a large database of temporal lobe seizures to investigate changes in intracranial EEG power during focal preserved and focal impaired consciousness seizures. In accordance with previously reported findings, we demonstrate that impaired consciousness is related to distinct electrophysiological changes in the temporal and extra-temporal lobes during FIC and FPC seizures. FIC seizures had significantly larger power than FPC seizures, with prominent increase in beta power in the lateral temporal (LT) lobe and delta power in the extra-temporal (ET) regions. Moreover, we found that a combination of these features of intracranial EEG power could discriminate FIC from FPC seizures with up to 86% accuracy supporting the idea that impaired consciousness in TLE is associated with the spread of seizure activity outside of mesial temporal lobe and slow wave (1-3 Hz) activity in the neocortex. We also conducted an in-depth analysis of seizures to address the three long-standing questions in the field (Figure 4 and Figure S 6). We confirmed the previous notion that bilateral seizures are more likely to cause impairment than unilateral seizures and neocortical slowing predicts impairment during TLE seizures. Contrary to previous notion, we didn’t find a relationship between onset side of seizure (left vs. right temporal lobe) and probability of impairment. Both left and right sided seizures could impair or preserve consciousness.

According to previous studies (Bancaud et al., 1984; Gloor, 1980; Inoue & Mihara, 1998; Jasper, 1964; Munari, 1980), occurrence of impaired consciousness during TLE seizures is related to bilateral involvement of temporal lobes. Along these lines, three other studies (Campora et al., 2024; Inoue & Mihara, 1998; Lux et al., 2002) report a higher occurrence of impaired consciousness in bilateral temporal seizures. Inoue and Mihara reported a correlation between impaired consciousness and propagation of seizure activity to contralateral temporal lobe with significantly higher probability (90% vs. 54%) of impaired responsiveness in patients with bilateral seizures compared to patients with unilateral seizures. Lux similarly found that bilateral temporal seizures impaired consciousness more frequently than left or right sided seizures (Lux et al., 2002). In a smaller study, Campora and colleagues reported a similar trend (non-significant) wherein 100% of bilateral seizures had loss of consciousness compared to 43% of unilateral seizures (Campora et al., 2024). Our study results resonate with these previous findings and confirm that bilateral seizures indeed had a significantly higher probability of impairing consciousness than unilateral seizures, but bilateral involvement is not a necessary condition for impairment. Seizures that don’t spread to the contralateral temporal lobe can still affect consciousness as is seen in 37 % of unilateral seizures in our study.

Unlike the above, consensus hasn’t been achieved on the effect of lateralization (left vs. right) on consciousness during temporal lobe seizures. Despite multiple studies trying to address this question, we observed two possible reasons for the disparity in results, i) small sample size and unavailability of bilateral temporal implants to determine true lateralization and spread of seizures and, ii) failure to recognize effect of seizure onset in language dominant vs non-dominant hemisphere. Most of the behavioral assessments of consciousness performed during seizures rely on speech and language functions which are impacted when seizures originate on the language dominant hemisphere causing aphasia, amnesia or speech arrest (Gabr, Luders, Dinner, Morris, & Wyllie, 1989; Marks & Laxer, 1998). If not carefully accounted for, these assessments may lead to conclusions that temporal lobe seizures originating on left side are more likely to cause impaired consciousness than seizures beginning on right side. While some studies (including ours) include non-verbal responsiveness and ability to follow commands in the assessment to mitigate the bias, it is crucial to differentiate seizures not just by the hemisphere of onset but also by language dominance to capture the full picture. We identified 12 categories based on seizure onset (left vs. right) and language dominance (dominant vs. non-dominant hemisphere of onset) for seizures with bilateral (mesial temporal) implants only, ipsilateral implants only and any implants (Figure S 6). In patients with bilateral implants, occurrence of impairment was not significantly different (p>0.05) for left vs. right seizures and for seizures originating in language dominant vs. non-dominant side. Similarly, there was no apparent relationship between impaired behavior and onset side or language dominance onset for seizures with ipsilateral implants only. However, there happened to be more left sided seizures for this group and more right sided seizures in bilateral implant group which then cause a false significant result when combining seizures with ipsilateral only and bilateral implants. Overall, our results show that both seizures originating on the left and right temporal lobe or language dominant and non-dominant hemisphere have the potential to impact consciousness. We also find that it is not sufficient to predict impairment based on whether one or both temporal lobes are involved in seizure but also requires information about the involvement of extra-temporal regions such as thalamus and neocortex.

Among other mechanisms of impaired consciousness, network-inhibition hypothesis postulates that temporal lobe seizures cause impairment when seizure activity extends beyond temporal lobe reaching the subcortical arousal systems including thalamus which then leads to depressed arousal and sleep like slow-wave activity in the frontal-parietal association cortices (Englot et al., 2010; Norden & Blumenfeld, 2002; Yu & Blumenfeld, 2009). While our group has previously implicated the role of cortical slowing in FIC seizures, in this study we additionally show the relative importance of temporal and extra-temporal intracranial EEG features in predicting impaired consciousness for a given seizure. Extra-temporal delta or neocortical slow wave activity and lateral temporal beta were the best predictors of impairment in the ictal period while mesial temporal beta activity was the least useful single predictor of impairment. A combination of ipsilateral and contralateral features of temporal and extra-temporal features gave the best classification accuracy suggesting that the pattern of seizure activity and its propagation outside the temporal lobe either ipsilaterally or bilaterally are key pieces to identifying if consciousness will be impacted during a seizure.

Despite a reasonably large sample size (186 seizures from 51 patients), we address a few limitations of our study. First, although we had 25 patients with bilateral implants in the mesial temporal lobe, our dataset didn’t have enough bilateral seizures of both types (FIC and FPC) to conduct within patient comparisons. Similarly, it was not possible to do within patient comparison of the effect of onset side on impairment during strictly unilateral seizures as only 4 of 25 patients had unilateral seizures originating from both hemispheres. Future studies should aim to create datasets consisting of more patients with bilateral implants with heterogeneous seizures (by onset side and type) per patient to allow for both within and between patient analysis. We think that having this type of dataset will further allow more specialized analysis of intracranial EEG such as synchrony between limbic and cortical regions for bilateral or unilateral seizures with and without impaired consciousness and shed light on why some bilateral seizures may preserve consciousness while some unilateral seizures cause impairment.

To summarize, we provide evidence through intracranial EEG analysis that on average FIC seizures exhibit two types of characteristics, fast beta activity in the ipsilateral temporal lobe (specifically lateral temporal) and bilateral slow wave delta activity in the frontal-parietal association cortices. Through a simple machine learning model, we demonstrated the predictive strength of these electrophysiological features for impaired consciousness in mesial temporal lobe seizures. Lastly, we established that the relationship between lateralization of seizure and probability of impairment is complex and there may be different mechanisms through which a unilateral or bilateral seizure or left vs right onset seizure affect consciousness.

## Supplementary figures

**Figure S 1.**
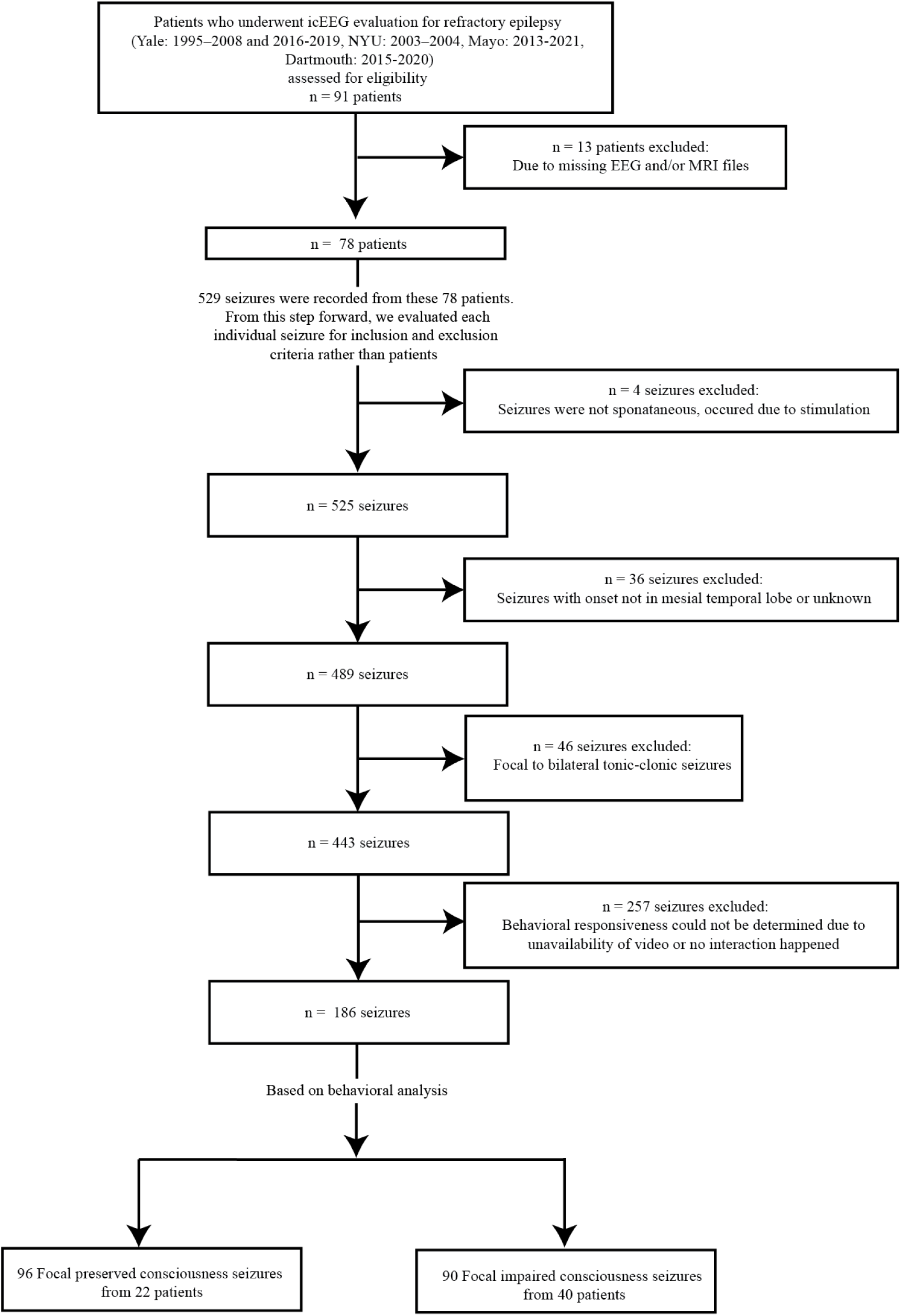
Consort diagram

**Figure S 2.**
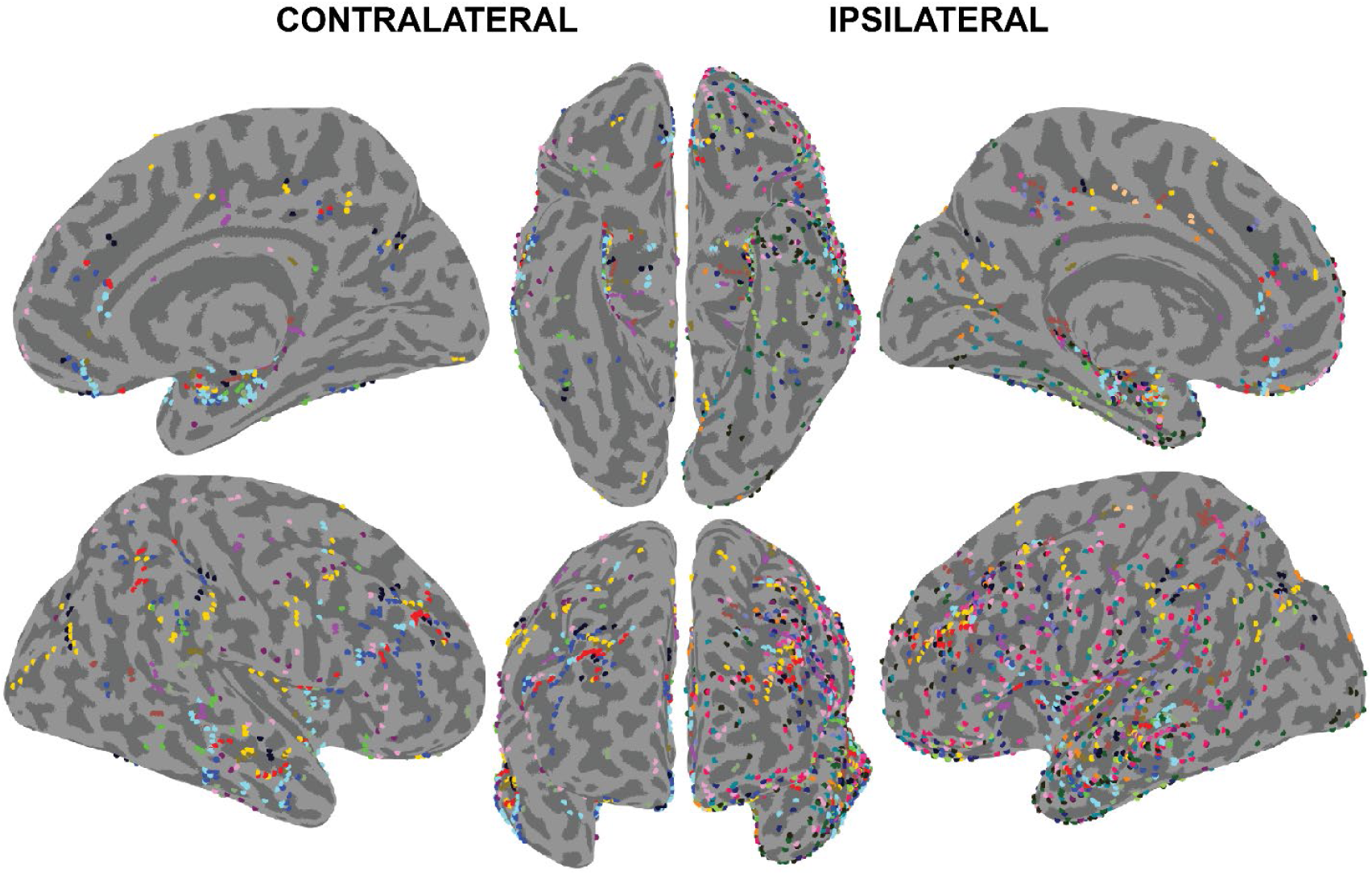
Electrode distribution in 23 patients for ipsilateral and contralateral brain regions (relative to seizure onset side). Lateral, medial, ventral and anterior views of both hemispheres are shown on an inflated brain. Each patient’s gray matter electrodes are colored by a different color and displayed at the closet vertex. Total gray matter electrodes across patients = 3097 (2211 ipsilateral, 886 contralateral).

**Figure S 3.**
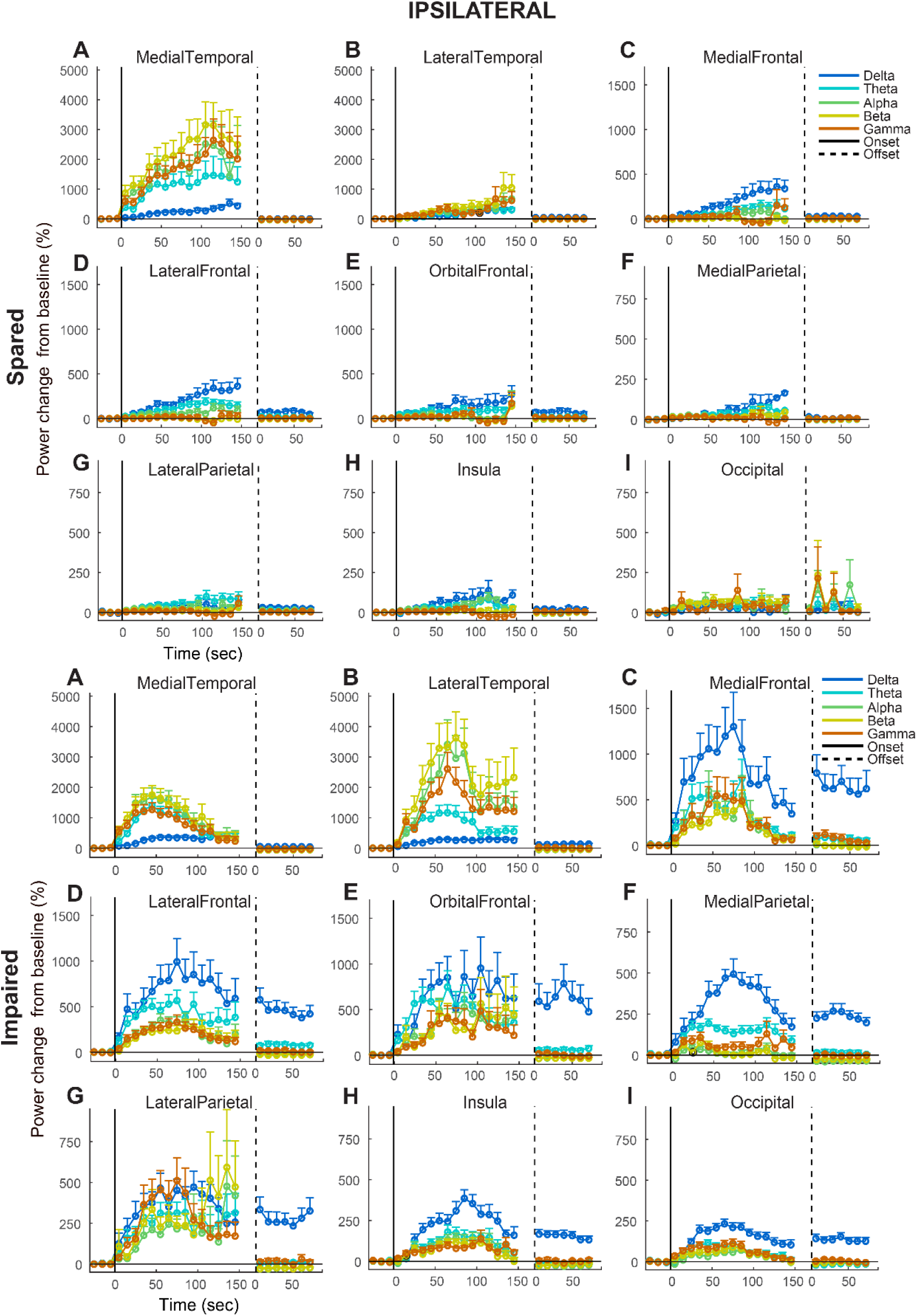
Bandpower time course plots of icEEG recorded from ipsilateral hemisphere during spared and impaired seizures across nine brain regions. Plots show percentage change in power during ictal and postictal periods relative to baseline. Solid vertical and dotted vertical lines indicate seizure onset and offset respectively.

**Figure S 4.**
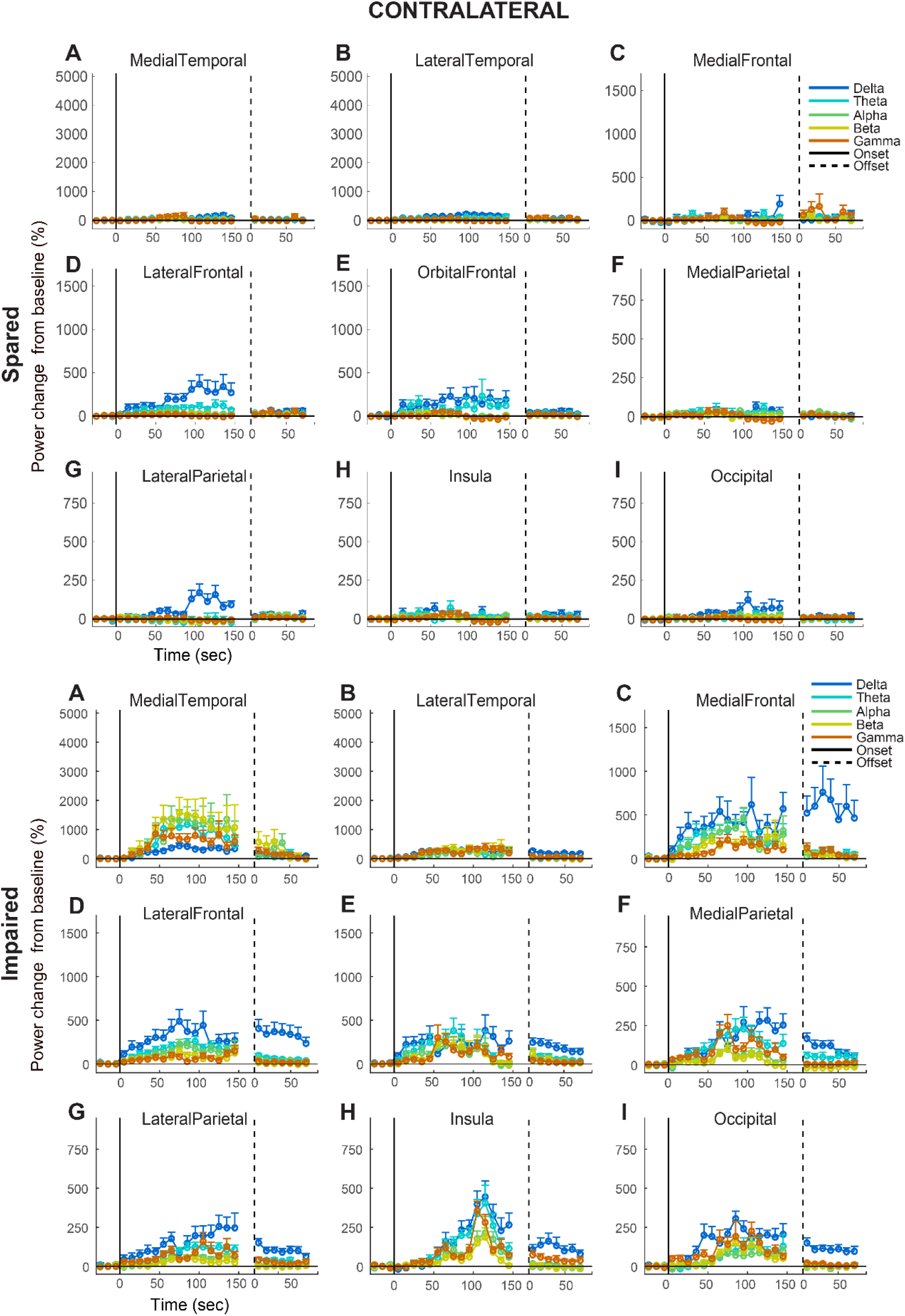
Bandpower time course plots of icEEG recorded from contralateral hemisphere during spared and impaired seizures across nine brain regions. Plots show percentage change in power during ictal and postictal periods relative to baseline. Solid vertical and dotted vertical lines indicate seizure onset and offset respectively.

**Figure S 5.**
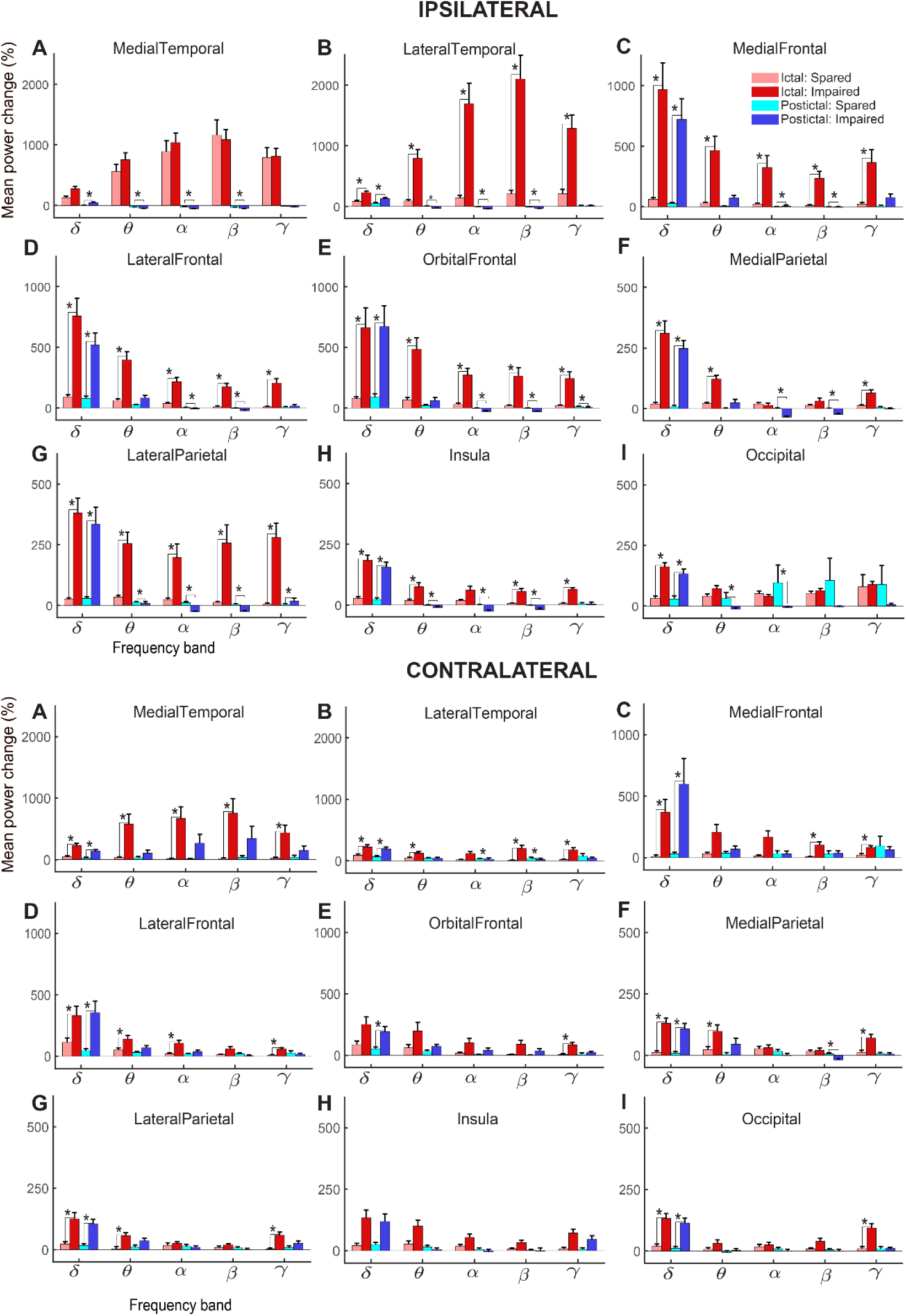
Bar plots show (+ SEM) percent change in icEEG power of different frequency bands (Delta, Theta, Alpha, Beta and Gamma) of spared and impaired seizures during ictal (pink and red) and postictal (cyan and blue) periods. Top plots show data from ipsilateral contacts across nine brain regions and bottom plots show the same for contralateral contacts. Note: different scale for y-axis across brain regions.

**Figure S 6.**
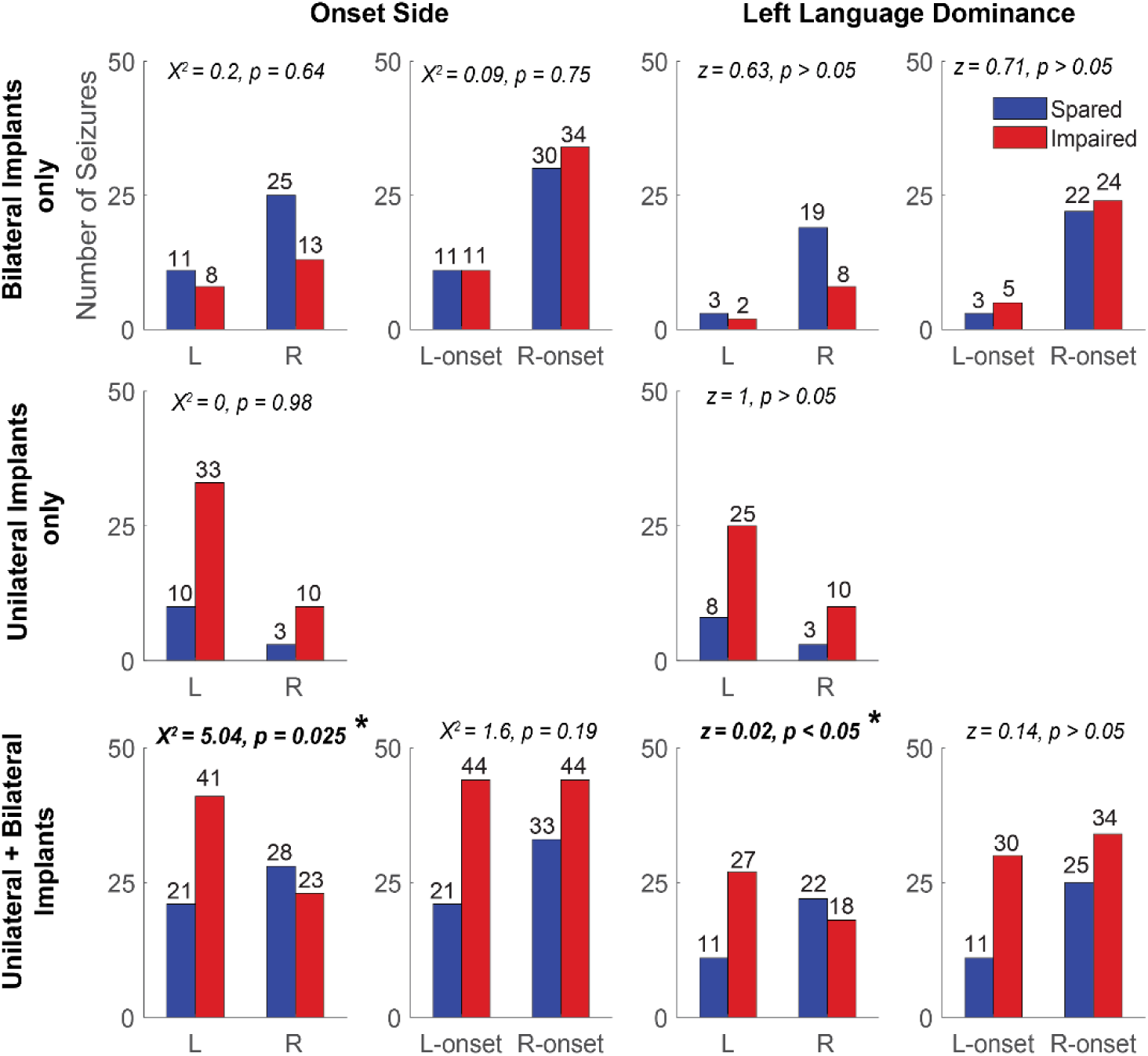
Distribution of seizures into left (L) or right (R) sided and left-onset or right-onset based on icEEG onset side and language dominance (L-dominant hemisphere, R-nondominant hemisphere). Top row includes only seizures with bilateral mesial temporal implants; middle row includes seizures with only ipsilateral implants and bottom row includes all seizures recorded unilaterally or bilaterally.

